# Adeno-associated virus serotype tropism and transduction in murine dorsal root ganglia following direct injection

**DOI:** 10.64898/2026.10.24.761581

**Authors:** Michael O’Donnell, Arjun Fontaine, John Caldwell, Richard Weir

**Affiliations:** Biomedical Engineering Department, University of Colorado – Denver | Anschutz Medical Campus, Aurora, CO 80045, USA; Department of Cell and Developmental Biology, University of Colorado – Denver | Anschutz Medical Campus, Aurora, CO 80045, USA; Rocky Mountain Regional VA Medical Center, Aurora, CO 80045, USA

**Keywords:** Anterograde Transport, Intraganglionic Injections, Mice, Optogenetics, Sensory Neurons

## Abstract

The peripheral nervous system (PNS) is highly heterogeneous, comprised of diverse subclasses of motor and sensory neurons. Optogenetics offers a high-resolution approach for interrogating neural circuits; however, optogenetic proteins must be selectively delivered to defined neuronal circuits in order to elucidate circuit-specific functional mechanisms. Adeno-associated viral vectors (AAVs) are widely used to deliver transgenes to the nervous system due to their relative safety and strong transgene expression. Multiple AAV serotypes are in common use, each capable of preferentially transducing distinct tissues and neuronal populations. However, characterization of the transduction behavior of these serotypes in the PNS, particularly in the dorsal root ganglia (DRG) and sensory nervous system (SNS), remain limited. This study evaluated the transduction efficiency of AAV8, AAV9, AAV2re, and PHP.S following direct intraganglionic injection into the DRG. AAV8, AAV9, and PHP.S exhibited comparable transduction efficiencies, whereas AAV2re transduced more than twofold fewer neurons. Despite similar overall efficiencies between AAV8, AAV9, and PHP.S, the neuronal volume distributions of transduced neurons were serotype-dependent. These results demonstrate that AAV serotype selection influences both the extent and size of transduced DRG neurons, indicating that serotype selection should be considered carefully when designing experiments that target peripheral sensory circuits.

## Introduction

The peripheral nervous system (PNS) plays a critical role in motion, sensory processing, and homeostasis (1, 2). The PNS is complex, broadly comprised of three distinct (motor, sensory, and autonomic) subdivisions, with each subdivision further comprised of multiple neuronal subclasses, each possessing unique transcriptional and electrophysiological properties (3–7). Due to its highly heterogeneous nature, the underlying mechanisms of PNS function remain incompletely understood and effectively treating its diverse pathologies is challenging. Given this complexity and limited understanding, there remains a critical need for methodologies to interrogate and modulate discrete peripheral neuronal circuitry.

Optogenetics uses gene editing approaches to create light-responsive tissues through the introduction of optical proteins into target cells. Utilizing optogenetic techniques in the nervous system provides a method to precisely monitor and control neural activity, both in ex-vivo environments and in freely moving animals (8–10). Through the introduction of various optical proteins into neural systems, nerve activity can be selectively stimulated or inhibited at a single-axon level while simultaneously monitoring neuronal signaling at an ensemble level (11–13). This level of resolution provides an extremely accurate method to map and study the PNS (14–16), modulate organ function innervated by the autonomic nervous system (17, 18), and investigate the neuronal circuits impacted by pathologies, particularly within the sensory nervous system (SNS) and its processes (19–22). However, to effectively utilize optogenetics to specifically modulate desired PNS processes, optical proteins must be precisely delivered to the target neuron population.

Adeno-associated viruses (AAVs) are non-enveloped viral vectors containing genetic constructs designed to express genes of interest in place of their native viral genome (23, 24). Widely used to deliver optogenetic constructs due to low immunogenicity and robust transgene expression, AAVs are also highly engineerable. Through strategic selection of appropriate promoter and serotype combinations, AAVs can be designed to restrict expression to desired tissues (25–27) and cell types (28–31). AAV serotypes exhibit distinct tissue tropisms based on capsid-receptor interactions at the cellular surface (32–34); consequently, identifying the optimal serotype for specific neuronal subtypes is critical for conducting effective SNS investigations. AAV8, AAV9, and AAV2-retro (hereafter called AAV2re) have all been shown to effectively transduce peripheral neurons through various routes of administration (35, 36), and AAV-PHP.S (hereafter called PHP.S) was engineered from AAV9 to more efficiently transduce the peripheral nervous system (37). The previously documented success of these serotypes in transducing PNS neurons makes them strong candidates for initial validation of serotype-specific transduction in the SNS.

An AAV’s transduction efficacy is influenced both by the serotype used to package the optogenetic protein genome as well as the route of administration used to deliver the AAV (38, 39). Localized injections yield higher localized expression of vector genomic copies per cell, enabling lower viral doses to achieve effective transduction while minimizing the risk of toxicity and immune responses (40–42). The dorsal root ganglia (DRGs) are the collections of peripheral afferent cell bodies that relay sensory information to the central nervous system (CNS) (43, 44), offering an ideal anatomical site to evaluate AAV serotype tropism and transduction efficiency in the SNS. Although all four chosen serotypes have been reported to transduce peripheral neurons, including DRG neurons in some cases, none have been evaluated following a direct DRG injection. In this study, we characterize the transduction efficacy and neuronal tropism of AAV8, AAV9, AAV2re, and PHP.S, with each driving EGFP transgene expression using the CAG ubiquitous promoter, following direct DRG injection into mice.

## Materials and Methods

### Animals

Adult male and female C57Bl/6J mice, aged 12-20 weeks at the time of injection, were used. Mice were ordered from Jackson laboratory. All experiments involving animals were conducted in accordance with the Institutional Animal Care and Use Committee of the University of Colorado – Denver | Anschutz Medical Campus.

### Viral Expression

All viral vectors were purchased from VectorBuilder (Chicago, IL, USA). Each vector expressed EGFP driven by the CAG promoter (AAV-CAG-EGFP-WPRE, 10^13^ GC/mL) and was packaged in one of the four selected serotypes (AAV8, AAV9, AAV2re, and PHP.S). Viral vectors were injected into murine L3 and L4 DRGs following a previously published methodology (45).

### Immunohistochemical Analysis

DRGs and sciatic nerves were excised and optically cleared following a previously published protocol (46). Post-clearing, DRGs were washed free-floating for 1 day in phosphate buffered saine with Triton (PBST) and then placed free-floating into primary antibodies (Goat anti-GFP, Abcam, 1:250 in PBST; Rabbit anti-NeuN, Abcam, 1:100 in PBST) for 5 days. Tissue was then washed in PBST for 1 day and placed, free-floating, in appropriate secondary antibodies (Alexa-647 conjugated donkey anti-goat, ThermoFisher, 1:250 in PBST; Alexa-568 conjugated donkey anti-rabbit, ThermoFisher, 1:250 in PBST) for 2 days. Tissue was washed a final time in PBST for 1 day and then washed overnight in 0.05M phosphate buffer (PB). Tissue was then mounted in VectaShield antifade mounting media (Vector Laboratories (H-1000-10)) on microscope slides with 240 µm spacers.

Spinal columns were excised, and the L2-L5 spinal segment was isolated and post-fixed overnight in paraformaldehyde (PFA, 4% in 0.1M PB) at 4°C. Additionally, plantar punches from the ipsilateral hind paw were collected using a 3mm biopsy punch tool and similarly post-fixed overnight. Tissue was removed from the fixative, placed in an EDTA solution (10% EDTA, 15% sucrose, 0.07% glycerol in 0.1M PB), and gently rocked overnight (plantar punches) or for 7 days (spinal segments) at 4°C. After decalcification, tissue was transferred to 30% sucrose in 0.1M PB and incubated with gentle rocking overnight (plantar punches) or for 7 days (spinal segments), before being mounted in optimal cutting temperature (OCT) compound and prepared for sectioning. Plantar punches were sectioned at 15 µm, while spinal segments were sectioned at 20 µm. Sections were mounted to microscope slides, placed on a slide warmer for 15 min at 40°C, and excess OCT was removed using a 0.05M PB wash. Plantar punch slides were then dried, and coverslips were mounted using Fluoromount-G Mounting Medium, with DAPI (ThermoFisher). Spinal section slides were placed tissue-side down on a staining platform and incubated with Nissl (NeuroTrace 640/660, ThermoFisher, 1:250 in PBS) for 4 hours at RT, before being similarly washed, dried, and coverslipped.

### Imaging

Fluorescent images of DRGs, sciatic nerves, and plantar punches were acquired using a 10x air, 20x air, or 40x oil objective on a Leica Stellaris 5 laser-scanning confocal microscope platform; composed of DMI8 microscope base, laser housing unit, LED epifluorescence box, and 5 HyD S multi-pixel photon counter detectors. Sequential multi-fluorescent images were collected using a 405 nm laser and tunable white light laser (WLL). Fluorescent images of spinal sections were acquired using an Olympus BX41 epifluorescence microscope using 4x air, 10x air, or 20x air objective with Semrock Filter sets (DAPI, FITC, TRITC, and Cy5).

Plantar punches were imaged using 405 nm laser and WLL tuned to 488 nm to observe DAPI staining and native EGFP fluorescence. DRGs and sciatic nerves were imaged with the WLL tuned to 568 nm and 647 nm to visualize fluorescent tags for appropriate antibodies. Images were acquired at 1024x1024 pixels and image stacks of the DRG 3-dimensional volume were obtained using Leica LASX software.

Spinal sections were imaged using the DAPI, FITC, and Cy5 filter sets on the Olympus to observe DAPI and Nissl staining alongside native EGFP fluorescence. Images were acquired and multi-color image overlay figures were generated using Olympus cellSens software and FIJI.

### Cell Quantification

DRG image stacks were analyzed using Imaris 10 “surfaces” tool. This tool allows users to both automatically and manually create surfaces encompassing fluorescent signal, allowing for identification of individual cells. Additionally, the tool calculates object statistics, such as volume and distance to other objects, to allow for broad physiological classification and colocalization analysis. Within the surface generation process, individual signal channels are selected, and a Gaussian filter is applied for background subtraction using a built-in algorithm. To calculate the lowest noise tolerance that would not identify positive cells, contralateral DRGs that were identically processed were used to calculate the lowest threshold that did not identify cells; observationally, a threshold value of 10 eliminated the majority of background autofluorescence in both the NeuN and GFP channels and precluded surface generation around neurons. This value was then applied to background subtraction, and surfaces were generated for all cells above said threshold.

Some resulting surfaces were composed of multiple neuronal cell bodies which had been merged together to form a single surface. All surfaces with a volume over 10,000 μm^3^ were manually examined to determine if they were comprised of multiple objects and multiple objects were manually separated. Additionally, any surface that was visibly merged (e.g., with sharp angle contours), regardless of volume, was manually separated as well. All GFP surfaces were evaluated for colocalization with NeuN, surfaces without any colocalization were manually evaluated. Surfaces that did not contain a neuron were eliminated; however, some GFP surfaces contained neurons with weak NeuN signal below the chosen threshold. For these neurons, a NeuN surface was manually generated. The quantity of labeled NeuN and GFP surfaces in individual DRGs was then averaged across serotypes.

Cell-volume size classes were extrapolated from thresholds reported in previous literature (47, 48). *Suzuki, et al.* classified neurons with a cross-sectional area less than 300 µm^2^ as small, while neurons larger than 600 µm^2^ were classified as large. In contrast, *Barabas, et al.* described neurons with soma diameters less than 27 µm as small and neurons with a soma diameter greater than 27 µm as large. A circular soma with a diameter of 27 µm has a cross-sectional area of ∼573 µm^2^, indicating that the small neurons defined by *Barabas, et al* includes both the small and medium populations defined by *Suzuki, et al.* Assuming spherical approximations to convert the published two-dimensional values into volume, these thresholds corresponded to cell volume boundaries less than ∼3900 µm^3^ for small neurons, and greater than ∼10,300 µm^3^ for large neurons. To account for possible variability in antibody labelling and volumetric reconstruction across individual cells, we designate neurons with volumes less than 4000 µm^3^ as small and neurons with volumes greater than 9000 µm^3^ as large.

## Results

### Quantification of viral transduction in the DRG

To evaluate the transduction efficiency of each serotype in DRG neurons, injected and contralateral DRG tissue was excised, optically cleared, and immunohistochemically stained prior to confocal imaging (Figure 1). Fluorescent labeling for both GFP (green) and NeuN (red) was observed in DRG neurons for all four serotypes (Figure 1A-D). Co-localization of the GFP+ signal was observed in subsets of NeuN+ cells across the imaging depth, supporting broad transduction of the DRG volume, and contralateral DRGs contained robust NeuN labeling while lacking GFP+ antibody expression (Figure 1E). The transduction efficiency of each virus was calculated by quantifying the number of NeuN+ cells that coexpressed GFP+ antibody staining in individual DRG volumes and averaging the efficiency across all DRGs per each serotype group (Figure 1F).

**Figure 1.**
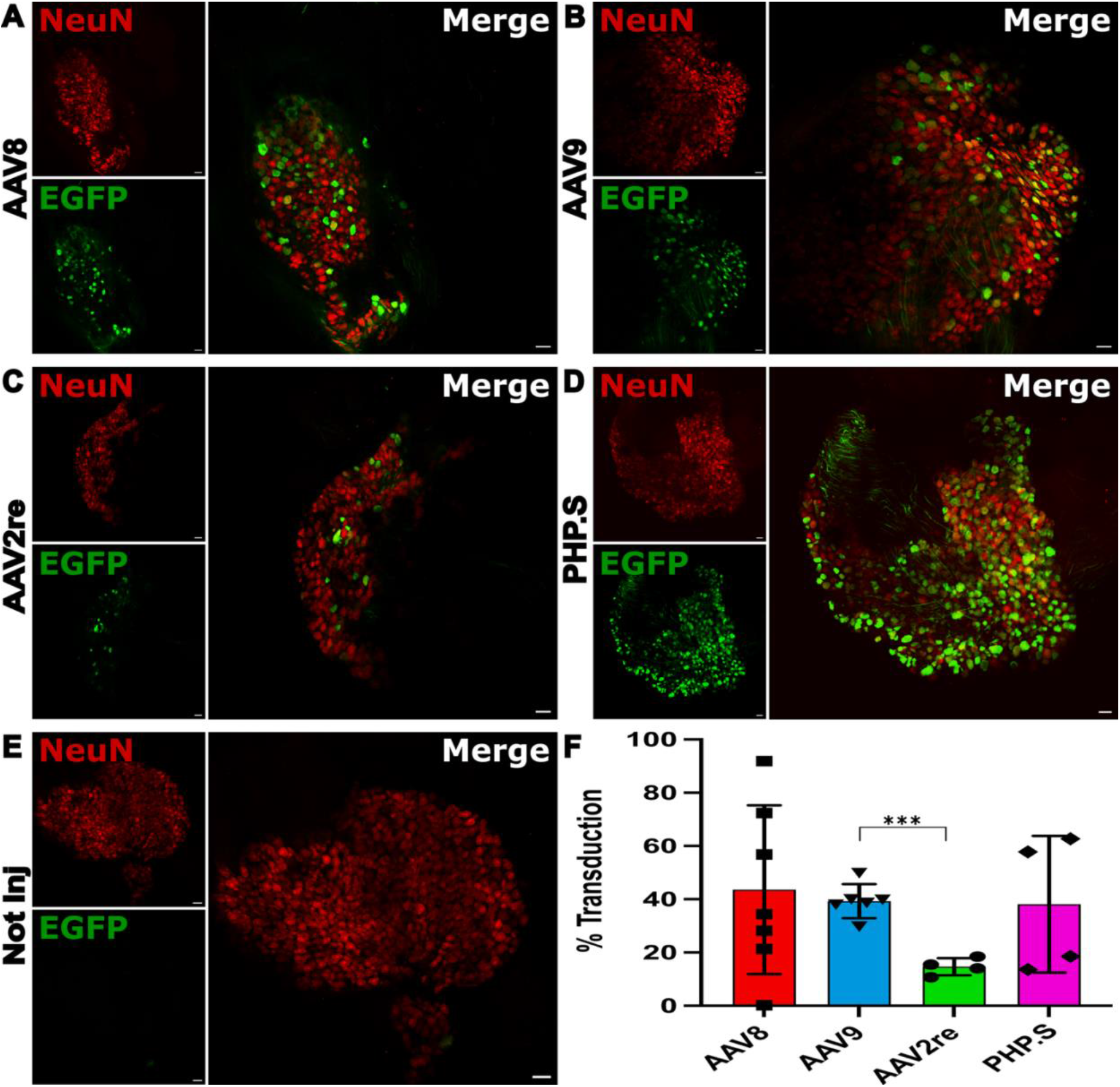
Intraganglionic injections of AAV8, AAV9, and PHP.S transduce DRG neurons at similar rates. Representative images of GFP expression in the DRG of AAV8-injected mice **(A)**, AAV9-injected mice **(B)**, AAV2re-injected mice **(C)**, PHP.S-injected mice **(D)**, and non-injected, contralateral DRG of injected mice **(E)**. Neurons marked with NeuN (scale bar, 50µm). **(F)** Quantification of transduced neurons in the DRG. Welch’s ANOVA indicated no significant difference between AAV8 (n = 7), AAV9 (n = 6), and PHP.S (n = 4) (p = 0.9421). Bars represent group means; error bars denote ±1SD.

To compare DRG-neuron transduction efficiency among serotypes, a one-way Welch’s ANOVA was performed to account for unequal variances between groups. Transduction efficiencies did not differ significantly among AAV8 (43.6 ± 31.7%), AAV9 (39.3 ± 6.4%), and PHP.S (38.2 ± 25.7%) groups (mean ± SD). In contrast, AAV2re transduced DRG neurons with more than two-fold lower efficiency (14.7 ± 3.2%). Dunnett’s T3 post hoc comparisons showed that AAV2re transduction efficiency was significantly different from AAV9 (adjusted p = 0.0002) but did not significantly differ from AAV8 or PHP.S (all adjusted p > 0.05). The lack of statistically significant differences between AAV2re and AAV8 or PHP.S likely reflects the greater variability observed in the AAV8 and PHP.S groups.

### Physiological classification of transduced neurons

The volumes from all GFP+ and NeuN+ cells across DRGs were collected for each serotype. Given that NeuN is a pan-neuronal marker, it can be assumed the cell volume distribution of the combined NeuN+ cells can approximate the physiological cell volume distribution of L3 and L4 murine DRGs. Cell volumes were plotted as histograms with 1000 µm^3^ bin widths, and each histogram was normalized to its maximum bin count. GFP+ distributions for each serotype were overlaid on top of the NeuN distribution to provide a direct comparison (Figure 2A). The distribution for NeuN+ cells was unimodal and right-skewed, as were the distributions for GFP+ cells from all serotypes. To provide visually descriptive summaries of these distributions, lognormal least square curves were fitted to each histogram. Dashed vertical lines indicate the peaks of the fitted curves, whereas diamonds mark the centers of the most populated histogram bins. Chi-square goodness-of-fit testing indicated departures from the fitted lognormal distributions; accordingly, the curves were used only as visual summaries.

**Figure 2.**
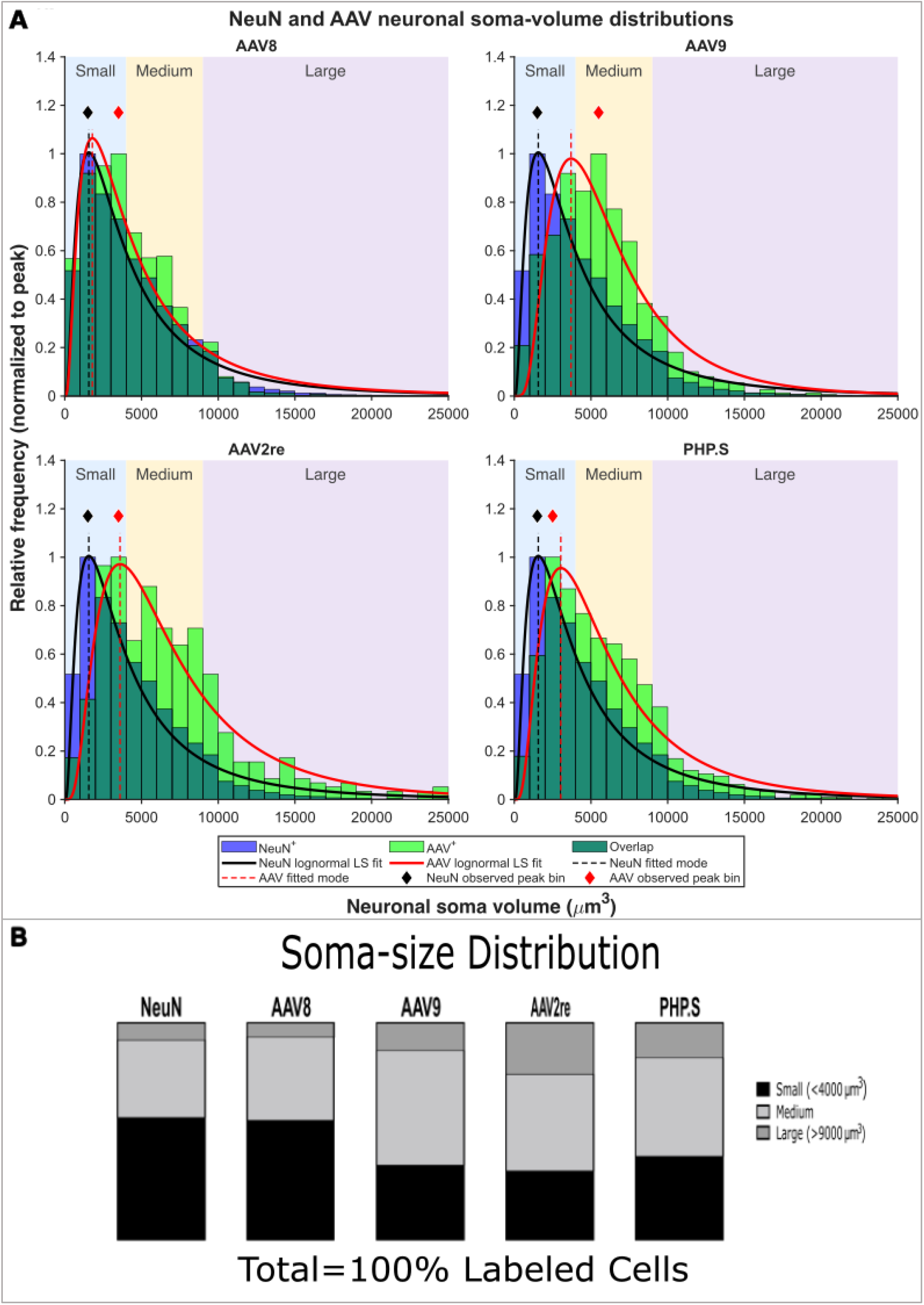
AAV9, AAV2re, and PHP.S transduce larger neurons compared to AAV8 and the native NeuN distribution. **(A)** Empirical soma-volume distributions normalized to maximum bin count (N_max_); bin width, 1000 µm^3^. The AAV8 distribution (N_max_ = 298) closely approximates the native NeuN population (N_max_ = 2682), while the AAV9 (N_max_ = 149), AAV2re (N_max_ = 58), and PHP.S (N_max_ = 207) distributions are shifted towards larger neurons. Solid lines show lognormal curves fit by least squares for visual comparison only. Dashed lines represent peaks of the lognormal curves, and diamonds indicate the most populated bins. **(B)** Size classification of transduced neurons; small (s), medium (m), and large (l). AAV8 (s: 54.94, m: 38.3, l: 6.76; %) follows a size distribution close to physiological approximation NeuN (s: 56.08, m: 35.57, l: 8.35; %), while AAV9 (s: 34.34, m: 52.57, l: 13.09; %), AAV2re (s: 31.62, m: 44.45, l: 23.93; %), and PHP.S (s: 38.28, m: 45.35, l: 16.37; %) transduce more medium to large neurons.

Visual comparison of the fitted curves shows that AAV8 most closely resembles the NeuN reference distribution, whereas AAV9, AAV2re, and PHP.S exhibit rightwards shifts towards larger soma volumes. The empirical peak-bin markers for all four serotypes are shifted toward larger volumes than the NeuN distribution, however, their locations do not consistently reflect the broader features of the fitted distributions. For example, the most populous bins for AAV8 and AAV2re are both within the 3000-4000 µm^3^ range, yet their fitted profiles differ substantially. The AAV8 fit peaks near the NeuN fit peak and follows a similar shape, while the AAV2re fit exhibits the most extended right-hand tail and peaks at a volume ∼2000 µm^3^ larger than NeuN. Similarly, AAV9 has the rightmost empirical peak bin, despite having a less extended tail than AAV2re and a similar fit peak location. These observations indicate that fitted profiles and empirical peak bins capture distinct, yet complementary, features of the soma-volume distributions; taken together, both may inform vector selection when targeting particular neuronal size ranges.

Statistical comparisons were performed on individual, unbinned soma-volume measurements. To compare soma-volume distributions between groups, a Kruskal-Wallis analysis was performed. The volume distributions differed significantly (p < 0.0001) between groups overall, indicating at least one group differed in distribution relative to the others. A Dunn’s post-hoc test revealed that volume distributions did not significantly differ between AAV8 and NeuN (p > 0.9999), nor AAV9 and PHP.S (p > 0.9999), while all other groups differed significantly (all adjusted p < 0.05). An additional Welch’s/Brown-Forsythe ANOVA with Games-Howell post hoc (for unequal variances) was performed as a sensitivity analysis. All groups differed significantly (all adjusted p < 0.05), excepting AAV9 and PHP.S (p = 0.7155). The concordant significance summary between Dunn’s and Games-Howell for the majority of the pairwise comparisons indicates that AAV9 and PHP.S do not significantly differ in cell volume distribution, while all other pairwise comparisons, excepting AAV8 and NeuN, have significantly different distributions. The AAV8 and NeuN pair comparison being significantly different under Games-Howell (p = 0.0324) while not different under Dunn’s (p > 0.999), signifies that the mean volume of AAV8 differs from NeuN, though the rank distribution does not.

Once it was confirmed that the majority of the tested serotypes transduced heterogenous distributions of neuronal volumes, the relative size distribution for each serotype was categorized. Primary sensory neurons can be broadly classified according to soma size (49, 50), with nociceptors typically classified as small, mechanoreceptors as medium-to-large, and proprioceptors as large. Accordingly, the size distribution of transduced neurons by each serotype could be an important factor in selecting the appropriate AAV serotype for somatosensory studies. Neuronal size classes are typically defined using diameter (µm) or cross-sectional area (µm^2^); here, a spherical approximation was used to convert measured soma volume into size classes, defining small neurons as less than 4000 µm^3^, medium neurons as 4000-9000 µm^3^, and large neurons as greater than 9000 µm^3^. AAV8 exhibited a size distribution similar to NeuN, approximating the physiological distribution of L3 and L4 murine DRGs, whereas AAV9, AAV2re, and PHP.S transduce a larger proportion of medium to large cells (Figure 2B). The size distributions suggest that AAV8 would be an ideal serotype for transducing smaller diameter neurons, such as nociceptors, while the other three serotypes would be more ideal for medium-to-large neurons, such as low-threshold mechanoreceptors (LTMRs) or proprioceptors, with AAV2re exhibiting the greatest bias towards large neurons of all four serotypes.

### Anterograde transduction along axons

DRG neurons are pseudo-unipolar with two bifurcating axonal branches, one innervating sensory end organs in the periphery and the other projecting into the CNS (spinal cord and brainstem). Direct intraganglionic injection into the DRG has been shown to transduce neuronal cell bodies without spread to other ganglia (51), and can induce expression in both plantar skin and the spinal column with serotype-dependent efficiency (52). The direction and magnitude of transduction away from the soma achieved by each serotype is therefore an important consideration in PNS modulation research. For the purposes of this manuscript, central anterograde transport is defined as transport from the soma to the spinal cord and CNS, while peripheral anterograde transport refers to transport to sensory end organs; in this specific case, plantar skin. Sciatic nerves were excised, optically cleared, and immunohistochemically stained and evaluated for GFP antibody expression. Additionally, plantar punches and the L2-L5 spinal segments were acquired from injected mice. Samples were cryosectioned and stained with DAPI and/or Nissl, then evaluated for observable EGFP expression.

Injected mice from all four serotypes contained GFP+ axons in ipsilateral sciatic nerves (Figure 3), indicating that neuronal expression in the DRG soma corresponds to expression further in the periphery. However, while AAV8, AAV9, and PHP.S successfully expressed EGFP in plantar sections (Figure 4), AAV2re lacked expression. Concurrent positive expression in both the sciatic nerve and plantar skin confirms that AAV8, AAV9, and PHP.S robustly transduce neurons and transport delivered proteins in a peripheral anterograde direction. The lack of expression in plantar skin for AAV2re, however, suggests it may not transport expressed proteins significantly in a peripheral anterograde fashion; it is possible that weak EGFP expression levels in sciatic axons may have been amplified by antibody staining while natural EGFP expression in plantar skin remained unobservable. Alternatively, due to the transduction bias towards larger neurons (i.e., proprioceptors) by AAV2re, it is possible that the end organs of transduced sciatic neurons do not project to plantar skin and rather terminate elsewhere in the periphery.

**Figure 3.**
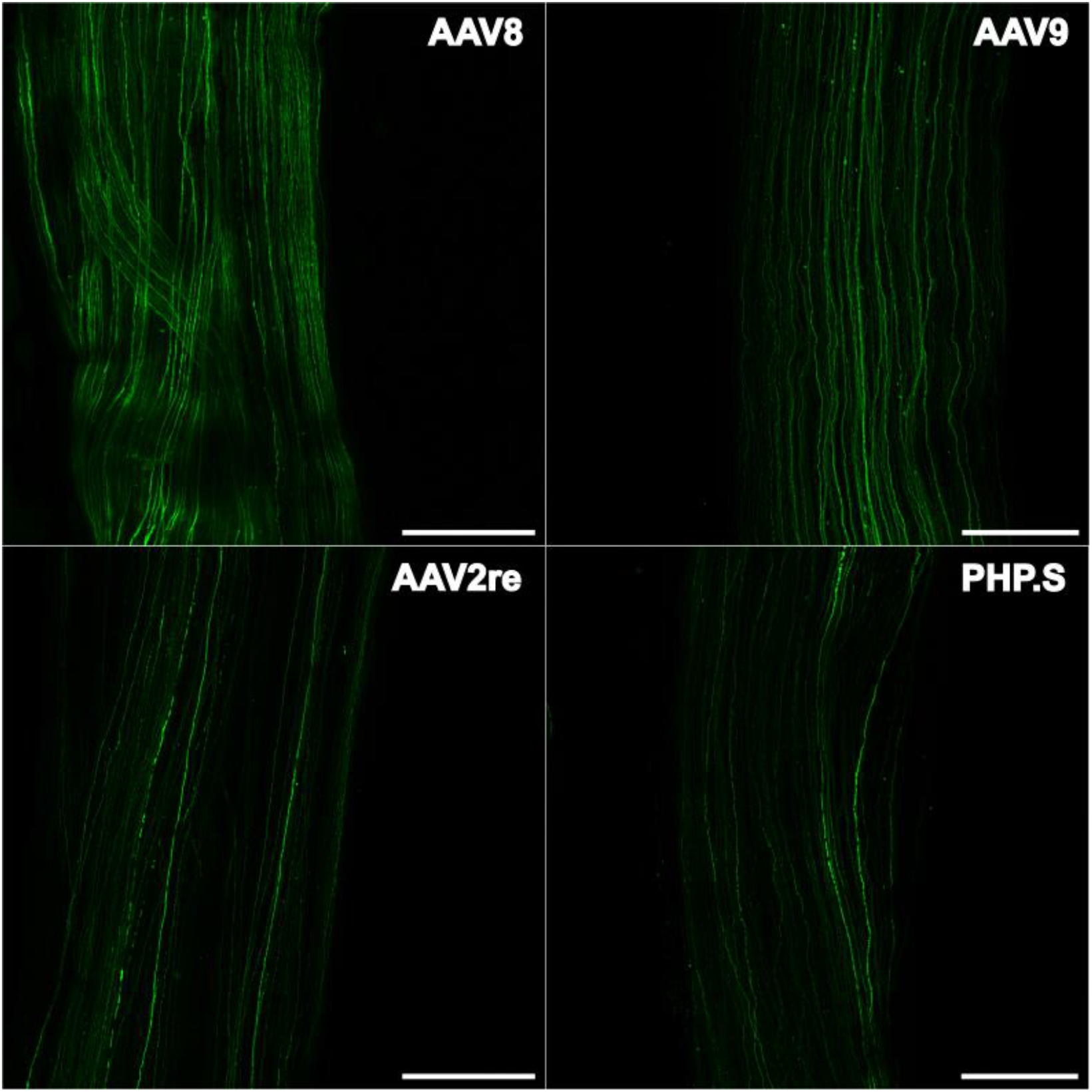
Direct DRG injection transduces sciatic nerve axons for all serotypes. Representative images of ipsilateral sciatic nerves demonstrate GFP expression (green) in axons for AAV8, AAV9, AAV2re, and PHP.S (scale bar, 250 µm)

**Figure 4.**
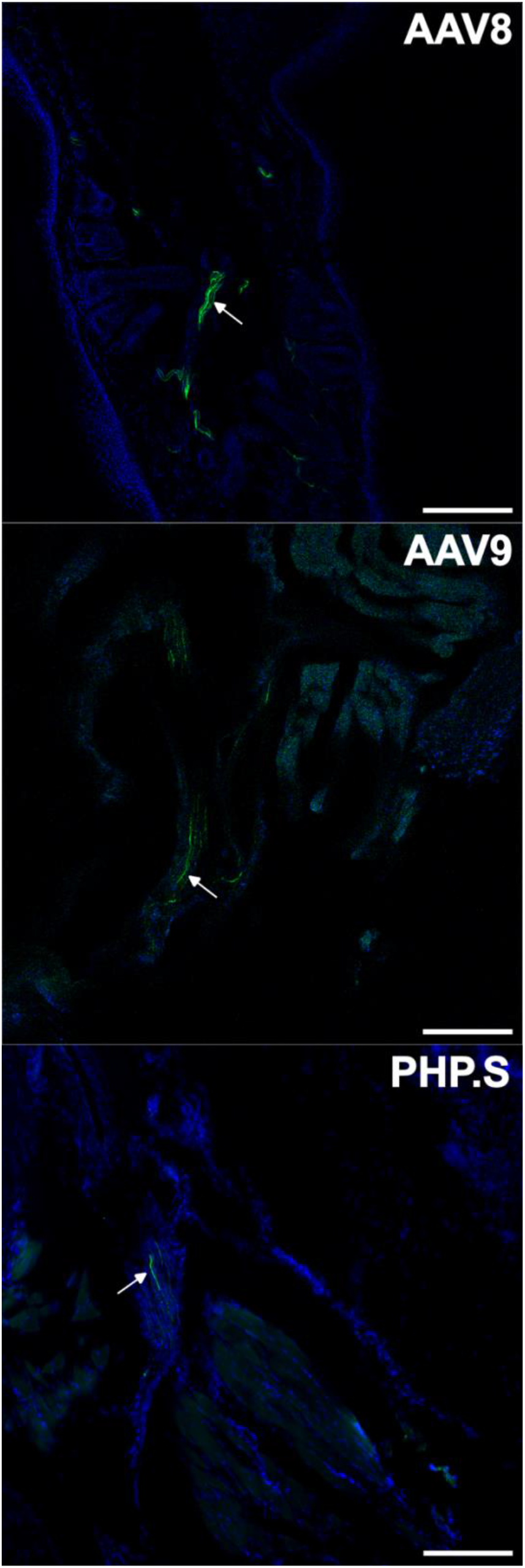
EGFP expression is present in plantar skin for AAV8, AAV9, and PHP.S. Representative plantar punch sections demonstrates EGFP expression (green) in axons present in plantar skin for AAV8, AAV9, and PHP.S. Tissue presence is confirmed with DAPI (blue) (scale bar, 200 µm).

Surgical complications during experimentation led to limited sample availability in central anterograde transport evaluation, as tissue samples were prioritized for DRG transduction quantification. AAV2re was engineered for enhanced retrograde transport (38, 53, 54) under various forms of administration and did not exhibit strong peripheral anterograde transport under the present injection paradigm. Additionally, PHP.S was modified from AAV9 (37); because PHP.S and AAV9 have significantly similar transduction efficiency, soma volume distribution, and peripheral anterograde transport behavior under the current experimental conditions, it was assumed that the central anterograde transport dynamic would likewise be similar. As such, AAV8 and AAV9 were chosen to be evaluated for central anterograde transport.

EGFP expression was observed in the dorsal root ganglia and dorsal root for both AAV8 (Figure 5 A,B) and AAV9 (Figure 6A). However, EGFP expression in the spinal column was only observed in the AAV9-injected spine (Figure 6B). Previous literature has demonstrated AAV8 expression in the dorsal horn of the spinal column following IM injection (35), indicating AAV8 is capable of central anterograde transport following retrograde transmission in sensory neurons. Additionally, positive expression in the dorsal root (Figure 5 A,B) may also indicate central anterograde transport capability following DRG injection, but lack of expression in the spinal column precludes a concrete conclusion. Given the limited sample availability for central anterograde evaluation (n = 1), it is possible that EGFP expression was present but spatially restricted and not represented in the sections obtained during cryosectioning.

**Figure 5.**
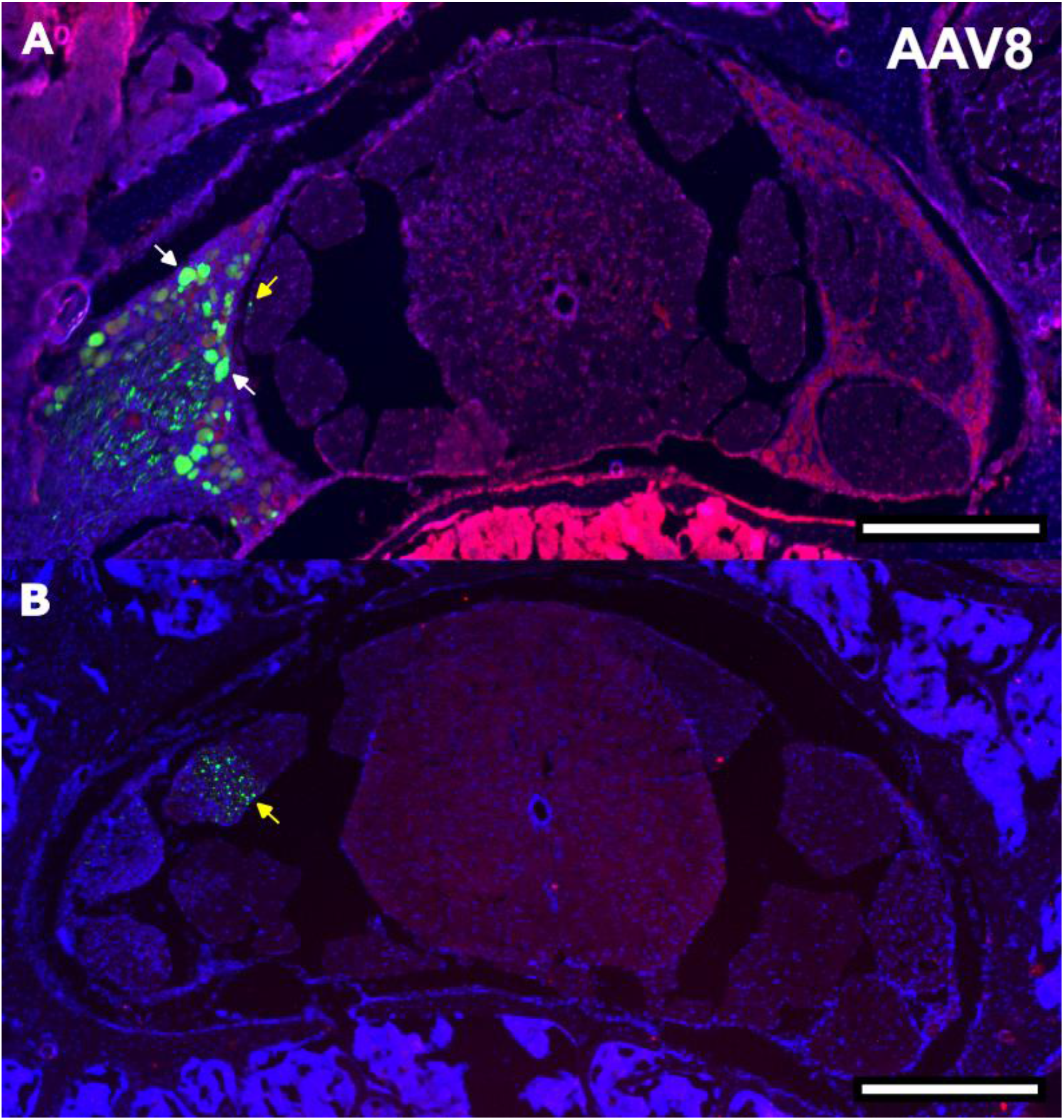
AAV8 expresses GFP in DRG and dorsal roots. **(A)** Representative image of spinal column demonstrates GFP (green) expression in DRG somata (white arrows), colocalized with Nissl (red), and axons of the dorsal root (yellow arrow) (scale bar, 500µm). **(B)** GFP expression in dorsal root (yellow arrow) without DRG present in section (scale bar, 500 µm). DAPI (blue) marks central canal and nuclei present in section.

**Figure 6.**
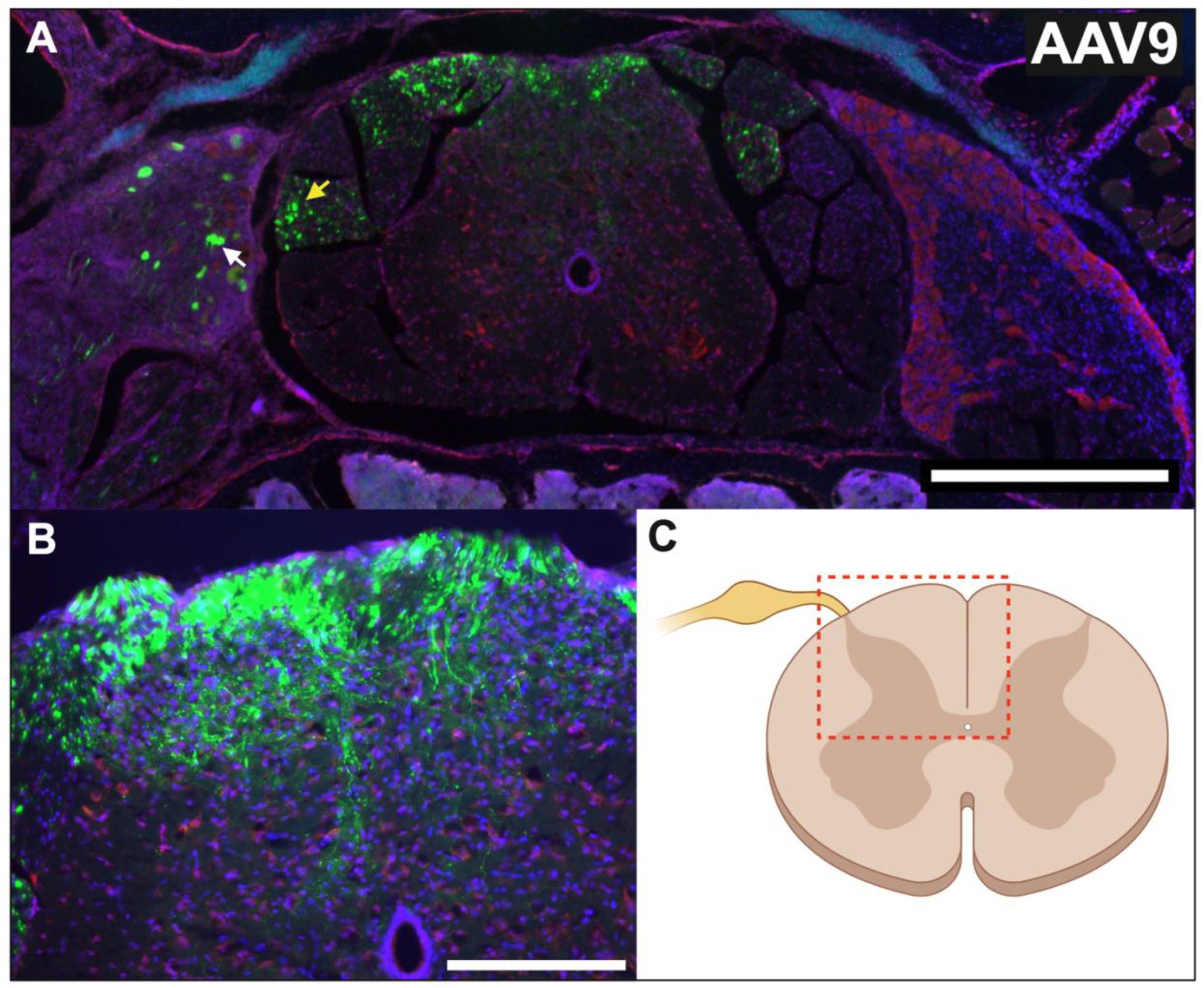
AAV9 successfully transduces axons in a central anterograde fashion following direct DRG injection. **(A)** GFP (green) expression in DRG somata (white arrow), dorsal root (yellow arrow), and dorsal horn of spinal cord (scale bar, 500 µm). **(B)** GFP expression extends deep into multiple laminae of dorsal horn, toward the central canal, identified by colocalization of DAPI (blue) and Nissl (red) (scale bar, 200 µm). **(C)** Representative schematic of transverse spinal section at the L3 level. Red, dashed box is representative of section image in (B). Created in BioRender. O’Donnell, M. (2026) https://BioRender.com/4zj3jlr

Contralateral EGFP expression can be observed in the AAV9 spinal column and dorsal roots (Figure 6A). Intraganglionic injection into the DRG typically restricts expression to the injected ganglia (51), so this expression is most likely caused by viral leakage into the intrathecal cavity from a substandard injection. GFP expression is not observed in the majority of observed contralateral DRGs, however, only one spine was evaluated for central anterograde transport. Further work is needed to determine whether the contralateral expression is truly a surgical anomaly.

## Discussion

The PNS is a highly complex system, comprised of diverse neuronal subclasses with distinct transcriptional and electrophysiological properties that discretely contribute to motor control and sensation. The large degree of heterogeneity in the PNS complicates efforts to resolve the functional organization of PNS neural circuits and contributes to the difficulty of developing effective treatments for PNS pathologies. Consequently, addressing these challenges requires higher-resolution methodologies to elucidate the functional mechanisms underlying sensory or motor responses. Optogenetics has revolutionized neuroscience research, allowing researchers to utilize light to stimulate or inhibit nerves at a single axon level while simultaneously monitoring ensemble neuronal activity.

However, the heterogeneity of the PNS requires optogenetic proteins to be delivered to defined neuronal populations. Achieving this specificity requires targeting strategies capable of restricting opsin expression to relevant neuronal subtypes at a genetic level.

AAVs have been widely adopted for targeted gene delivery to specific neuronal populations due to their low immunogenicity, robust expression, and serotype-dependent tropism. AAV serotype capsids determine their interactions with cell-surface receptors, thereby influencing tissue specificity and transduction efficiency. This work compares the transduction efficiency and cellular tropism of four AAV serotypes following direct DRG injection. Although AAV8, AAV9, AAV2re, and PHP.S have each been reported to effectively transduce peripheral neurons, their performance has never been directly compared following direct intraganglionic injection.

No significant difference was observed in the transduction efficiency of AAV8, AAV9, and PHP.S, with all three serotypes transducing approximately 40% of NeuN-labeled neurons within the DRG. AAV8 exhibited the largest variability within transduction efficiency, with one sample transducing ∼90% of available neurons, while another transduced less than 1% of available neurons. Although this low value may appear to be an outlier, no experimental anomalies or differences were observed and therefore there was no justification for its exclusion. In contrast, AAV2re only transduced ∼15% of DRG neurons. This finding contrasts with a previous study (55) that reported no appreciable difference in transduction efficiency of DRG neurons between AAV9, PHP.S, and AAV2re following systemic delivery in neonatal rats. This discrepancy could be caused by species- or age-related differences, as the present work injected adult mice, or may be attributable to difference in injection location. Due to the variability observed with AAV8 and PHP.S, the transduction efficiency of AAV2re was significantly different only from AAV9.

The volumes of all NeuN-labeled neurons across all experimental conditions were combined to approximate the physiological volume distribution of neurons in L3 and L4 DRGs. This distribution was unimodal and right-skewed, reflecting a predominance of small neurons (>50%). The volume distributions of GFP+ neurons for each serotype were also unimodal and right-skewed; however, only AAV8 was significantly similar to the native NeuN distribution (Kruskal-Wallis with Dunn’s post-hoc, p > 0.9999). The other three serotypes preferentially transduced medium to large neurons, consistent with previous literature (38, 55). AAV9 and PHP.S did not differ significantly in either their mean neuronal volume (Welch’s Anova with Games-Howell post-hoc, p = 0.7155) or in their rank distribution (Kruskal-Wallis with Dunn’s post-hoc, p > 0.9999), while AAV2re was skewed more towards large neurons and was significantly different (all adjusted p < 0.05) from all three other serotypes as well as the native NeuN distribution. This bias may provide an additional explanation for the lower overall transduction efficiency of AAV2re, as maximal efficiency of AAV uptake occurs near the injection site. Because AAV2re preferentially targets large neurons, and large NeuN-labeled neurons constitute a small fraction of the total DRG population, this tropism may limit the proportion of neurons that AAV2re can transduce within the DRG.

The differences in neuronal size distributions transduced by each serotype may provide a powerful tool for various SNS research. Nociceptive neurons are typically small and unmyelinated, whereas LTMRs and proprioceptors are larger and myelinated. Given that AAV8 closely follows the native distribution of NeuN-labeled cells and transduces a larger proportion of smaller neurons, it may be an ideal serotype for nociceptive studies. In contrast, because AAV9 and PHP.S preferentially transduce medium to large neurons, these two serotypes are likely better suited for studies involving LTMR and proprioceptors.

EGFP expression was observed in ipsilateral sciatic nerve axons for all four serotypes. To confirm the extent of peripheral anterograde transport, plantar punches were taken from ipsilateral hind paws, cryosectioned, and mounted with DAPI. EGFP expression was observed in plantar sections in AAV8, AAV9, and PHP.S. This indicates that these three serotypes will transport transgenes to the end-organs following DRG injection; this capability would allow researchers to stimulate nervous system circuits via the plantar skin or map specific sensory circuity in both the sciatic nerve and DRG following physiological stimuli. EGFP was not observed in AAV2re plantar sections, though positive expression in the sciatic nerve indicates partial anterograde transport. It is possible that antibody staining amplified weak expression levels in the nerve and native EGFP presence was too weak to be observed in plantar skin, though alternatively, due to the predisposition to transduce larger neurons such as proprioceptors, the AAV2re-transduced axons may have also terminated elsewhere in the periphery. More study will need to be conducted to evaluate if transduced axons end in hairy skin or muscle spindles, as only the glabrous hind paw skin was evaluated.

AAV8 and AAV9 were evaluated for central anterograde transport into the spinal column. EGFP expression was observed in dorsal roots for both serotypes, indicating successful anterograde transport; however, the AAV8 spinal sections lacked expression in the dorsal horn and spinal column. In contrast, AAV9 contained strong EGFP expression in the spinal column, extending through multiple laminae towards the central canal. Due to the small sample size (n = 1), the lack of expression in the spinal column for the AAV8 does not preclude successful anterograde transport to the spinal column, especially as previous work reports conserved mechanisms for anterograde transport between AAV8 and AAV9 in primary rat cortical neurons (56). Expression may have been spatially restricted and not represented in the collected sections.

The spinal sample for AAV9 contained EGFP expression on both the ipsilateral and contralateral sides of the spinal cord (Figure 6). Contralateral expression is not observed in most DRGs following intraganglionic injection, suggesting this expression is not a consistent feature. Because intraganglionic injection is expected to restrict expression to only the injected DRG (51), the contralateral signal most likely resulted from viral leakage from the DRG into intrathecal space, allowing exposure to contralateral tissue. Leakage can be caused by large injection volumes or from DRG damage caused by suboptimal injections. These observations suggest that future studies should use larger sample sizes to determine whether contralateral expression occurs as a consequence of surgical damage, while also evaluating transduction efficiency across different injection volumes to establish whether a smaller volume can achieve comparable results while minimizing the risk of leakage.

While these results demonstrate serotype-specific transduction patterns and transport following direct DRG injection, this study has certain limitations that should be considered. There was a limited availability of animals, leading to small sample sizes and preventing the evaluation of variability within groups and comparison of sex differences. Additionally, transduction was only assessed in the L3 and L4 DRGs of adult mice. While the majority of somatosensory studies have been conducted in rodents, there are species-specific molecular profiles and distributions that can also differ across DRG level (57). Understanding these differences will be vital for translating pre-clinical studies to clinical research. In this vein, neuronal classification was based entirely on soma size, which does not delineate function subtypes or molecular groups of sensory neurons. Finally, the occurrence of contralateral spinal expression suggests that minor leakage occurred during intraganglionic injection, contributing to off-target expression. To fully refine the tropism profiles of these serotypes following intraganglionic injection, future studies will need to employ larger sample sizes and evaluate multiple injection volumes and the molecular profiles of transduced neurons.

Characterizing the transduction efficiency and tropism of AAV serotypes across various routes of administration is only one component of achieving selective transgene delivery. Serotypes provide a broad tissue-level tropism, and viral expression can be further restricted through promoter selection (30, 58). Gene-specific promoters may restrict expression to defined neuronal populations (29, 59), enabling precision delivery of optogenetic tools to functionally distinct neuronal subtypes and circuits. Ongoing work in our laboratory uses epigenomic data to identify regulatory elements and engineer cell type-selective expression for preferential targeting of LTMRs, with initial results in preparation for publication. This approach could facilitate mechanistic investigation of diverse sensory circuits and, in the longer term, inform development of strategies for restoring sensory feedback, therapeutically modulating organ systems, and investigating neural pathologies.

In summary, all four serotypes are capable of transducing sensory neurons in the DRG following direct intraganglionic injection. AAV8, AAV9, and PHP.S did not differ significantly in transduction efficiency, yet they exhibited distinct neuronal size preferences. AAV8 transduces a larger proportion of smaller neurons, whereas AAV9 and PHP.S preferentially transduced medium to large neurons. These findings provide a foundation for selecting serotypes based on desired neuronal subpopulations. Further work will be needed to fully refine these transduction profiles across spinal levels and molecular neuronal subtypes; however, the present study represents an important step towards characterizing AAV tropism in the PNS and leveraging these tools for the targeted manipulation of sensory circuits.

## Data Availability

The data generated and analyzed during this study are available from the corresponding author on reasonable request.

## Disclosure of AI Usage

Perplexity AI was used solely to assist in generating code for the assembly and formatting of Figure 2a, which comprises four previously generated histograms. The code overlaid lognormal least-squares fit curves, placed a marker above the most populous bin in each histogram, shaded the background to delineate three size classes, and arranged the histograms into a 2 × 2 layout with a shared legend. Apart from fitting the lognormal curves, the AI-assisted code did not prepare, process, or analyze the underlying data. All AI-assisted code and the resulting figure panel were reviewed and verified by the authors. No other figures or figure panels were generated or modified using AI-assisted code.

